# DENTATE GRANULE CELLS ARE HYPEREXCITABLE IN THE TGF344-AD RAT MODEL OF ALZHEIMER’S DISEASE

**DOI:** 10.1101/2021.04.14.439848

**Authors:** Lindsey A. Smith, Anthoni M. Goodman, Lori L. McMahon

## Abstract

The dentate gyrus is both a critical gatekeeper for hippocampal signal processing and one of the first regions to dysfunction in Alzheimer’s disease (AD). Accordingly, the appropriate balance of excitation and inhibition through the dentate is a compelling target for mechanistic investigation and therapeutic intervention of early AD. Previously, we reported increased LTP magnitude at medial perforant path-dentate granule cell (MPP-DGC) synapses in slices from TgF344-AD rats compared to Wt as early as 6 months. We next determined that the enhanced LTP magnitude was due to heightened function of β-ARs leading us to ask if dentate granule cells (DGCs) also have modified passive or active membrane properties which may also contribute to hyperexcitability or aberrant hippocampal processing. Although there was no detectable difference in spine density and presynaptic release probability, dentate granule cells (DGCs) themselves might have increased electrical response to synaptic input during LTP induction, which was measured as a significant increase in charge transfer during high-frequency stimulation. In this study, we found passive membrane properties and active membrane properties are altered, leading to increased TgF344-AD DGC excitability. Specifically, TgF344-AD DGCs have an increased input resistance and decreased rheobase, decreased sag, and increased action potential (AP) spike accommodation. Importantly, we find that the voltage response increased in DGCs from TgF344-AD compared to Wt accompanied by decreased delay to fire the first AP, indicating that for the same amount of depolarizing current injection TgF344-AD DGCs membranes are more excitable. Taken together, DGCs of TgF344-AD rats are more excitable, but due to heightened accommodation, may be unable to discharge at high frequency for longer durations of time, compared to their Wt littermates.

## Introduction

Increased neuronal activity is directly linked to the production, secretion, and regional deposition of Aβ (Wei et al., 2010; Yamamoto et al., 2015) and tau (Wu et al., 2016; Vogel et al., 2020). While the locus coeruleus (LC) is now known to be the first site of pathological tau accumulation (Braak et al., 2011), cortical regions associated with high metabolic activity, such as the default mode network, including the entorhinal cortex and hippocampus are early targets of network abnormalities in Alzheimer’s disease (AD) (Reiman et al., 2012; Vogel et al., 2020). Functional imaging studies reveal hyperactivity in the hippocampus during mild cognitive impairment (Dickerson et al., 2004) and even in preclinical AD (Quiroz et al., 2010; Reiman et al., 2012). While decreased synaptic inhibition is commonly observed in AD-mouse models (Rocher et al., 2008; Verret et al., 2012; Hazra et al., 2013), compensatory remodeling of inhibitory circuits is thought to result from early aberrant excitation (Palop and Mucke, 2016). In support, APP23xPS45 mice have increased activity of hippocampal CA1 pyramidal neurons prior to plaque formation (Busche and Konnerth, 2016). Additionally, increased intrinsic excitability from mature DGCs is reported in Tg2576 AD mice (Nenov et al., 2015; Hazra et al., 2013).

We previously reported increased postsynaptic depolarization during LTP-inducing stimulation and enhanced LTP magnitude at medial performant path synapses onto dentate gyrus granule cells (MPP-DGC) in 6 month old TgF344-AD rats (Goodman et al., 2020; Smith and McMahon, 2018), a rodent model which more fully recapitulates AD-like pathology in an age-dependent manner (Rorabaugh et al., 2017; Smith and McMahon, 2018; Cohen et al., 2013; Tsai et al., 2014; Pentkowski et al., 2018; Voorhees et al., 2018; Muñoz-Moreno et al., 2018; Bazzigaluppi et al., 2018; Joo et al., 2017; Do Carmo and Cuello, 2013). Specifically, the spatiotemporal spread of synaptic dysfunction in hippocampus begins in the dentate gyrus (DG) prior to area CA1 in both male and female TgF344-AD rats. At 6 months of age, amyloid plaques are first detected in hippocampus, tau tangles are absent, but gliosis is significant (Cohen et al., 2013). In addition, we recently reported significant loss of noradrenergic fibers in hippocampus beginning at 6 months (Goodman et al., 2020) when accumulation of hyperphosphorylated tau (pTau) is present in the locus coeruleus (Rorabaugh et al., 2017), the origin of hippocampal noradrenergic (NA) innervation (Loy and Moore, 1979). Furthermore, concurrent with degeneration of hippocampal NA input, we observed heightened function of β-adrenergic receptors (β-ARs) at MPP-DGC synapses that were responsible for the enhanced LTP magnitude at 6-months in TgF344-AD rats (Goodman et al., 2020). Importantly, while the β-AR antagonist propranolol prevented the heightened LTP magnitude in TgF344-AD rats, it did not completely abolish the increased in steady state depolarization during the tetanus (Goodman et al., 2020), suggesting that another mechanism is contributing to the increased postsynaptic depolarization.

Heightened intrinsic excitability of DGCs could explain the increase in steady-state depolarization during the tetanus. Thus, we investigated this using whole-cell current clamp recordings of DGCs from TgF344-AD and Wt littermate females to assess passive and active membrane properties. Although some information exists showing increased excitability in the dentate of transgenic mice with the same transgenes (Hazra et al., 2013; Nenov et al., 2015), it is not known whether this also occurs in DGCs in the novel TgF344-AD rat model. Here, we report decreased rheobase, increased voltage response, increased probability of firing an AP, decreased sag voltage, and greater spike accommodation in 6-month DGCs in TgF344-AD rats compared to Wt. Together these data show that TgF344-AD DGCs are hyperexcitable and this gain of function may contribute to the enhanced depolarization during tetanus used to induce LTP we observed at 6 months of age (Goodman et al., 2020; Smith and McMahon, 2018).

## Methods

### Animals

All breeding and experimental procedures were approved by the University of Alabama at Birmingham Institutional Animal Care and Use Committee and follow the guidelines outlined by the National Institutes of Health. TgF344-AD males harboring the amyloid precursor protein Swedish (APP_swe_) and delta exon 9 mutant human presenilin-1 (PS1_∆E9_) transgenes were bred to wildtype (Wt) F344 females (Envigo, Indianapolis, IN), as done previously in our lab (Smith and McMahon, 2018). Rats were maintained under standard laboratory conditions (12 h light/dark cycle, lights off at 14:00 h, 22°C, 50% humidity, food (Harlan 2916; Teklad Diets, Madison, WI) and water ad libitum). Animals were housed using standard rat cages (7 in. (height) × 144 in^2^ (floor)). Female TgF344-AD and Wildtype (Wt) littermates were aged to 6-8 months for these experiments. Transgene incorporation was verified by polymerase chain reaction (PCR) as described previously (Smith and McMahon, 2018).

### Surgery

To control for cycling hormones, and to replicate previously used conditions (Smith and McMahon, 2018), all female rats were acutely ovariectomized. Briefly, TgF344-AD and Wt littermate female rats were bilaterally ovariectomized (OVX) under 2.5% isoflurane in 100% oxygen, using aseptic conditions. To account for different estrous cycle stages at the time of OVX, a 10-day minimum postoperative interval was used prior to experimentation, which allows for the depletion of endogenous ovarian hormones as previously published (Smith and McMahon, 2005, 2006; Smith et al., 2010).

### Hippocampal slice preparation

Animals were deeply anesthetized via inhalation anesthesia using isoflurane, rapidly decapitated, and brains removed. Coronal slices (400μm) from dorsal hippocampus were prepared using a Leica VT 1000A vibratome (Leica Microsystems Inc, Buffalo Grove, IL). To preserve neuronal health and limit excitotoxicity, slices were sectioned in low Na^+^, sucrose-substituted ice-cold artificial cerebrospinal fluid (aCSF) containing [in mM: NaCl 85; KCl 2.5; MgSO_4_ 4; CaCl_2_ 0.5; NaH_2_PO_4_ 1.25; NaHCO_3_ 25; glucose 25; sucrose 75 (saturated with 95% O_2_, 5%CO_2_, pH 7.4)]. Slices were held at room temperature for 1 h in HEPES buffered artificial cerebrospinal fluid (aCSF) [in mM: 92.0 NaCl, 2.5 KCl, 2.0 MgSO_4_, 2.0 CaCl_2_, 1.25 NaH_2_PO_4_, 30 NaHCO_3_, 25 Glucose, 5 L-ascorbic acid (saturated with 95% O_2_, 5%CO_2_, pH 7.4)] before transfer to submersion chamber for recordings. HEPES modified storing ringer was used to reduce cell swelling and slice damage and robustly improves slice health for path clamp recordings in aged animals (Ting et al., 2014). Following the 1 hr recovery, slices were transferred to a submersion chamber and continuously perfused (∼2-3 ml/min) with standard artificial cerebrospinal fluid (aCSF) [in mM: 119.0 NaCl, 2.5 KCl, 1.3 MgSO_4_, 2.5 CaCl_2_, 1.0 NaH_2_PO_4_, 26.0 NaHCO_3_, 11.0 Glucose (saturated with 95% O_2_, 5%CO_2_, pH 7.4)] held at 26–28 °C.

### Whole-Cell Current Clamp

Whole-cell patch clamp recordings were performed from dentate granule cells (DGC) in the ectal limb (upper blade). Somas of DGCs were blind patched using borosilicate glass pipettes (3-6 MΩ) filled with intracellular solution containing (in mM): 120 K-Gluconate; 0.6 EGTA; 5.0 MgCl_2_; 2.0 ATP; 0.3 GTP; 20.0 HEPES; 270-280 mOsm, pH 7.2 adjusted with KOH. To assess intrinsic excitability, somatic current injections were performed in the presence of glutamatergic antagonists, APV (100μM), RO 25-6981 [0.5μM], and DNQX (10μM), and the GABAergic antagonist Picrotoxin (100 μM), to block synaptic transmission. Nifedipine [10μM] was also used to block Ca_V1.1-4_ which contribute to heightened β-AR function in TgF344-AD rats (Goodman et al., 2020). After achieving whole-cell configuration, cells were voltage clamped at −70 mV for 3-5 minutes, and the amplifier was then switched to current clamp. Hyperpolarizing and depolarizing 800ms square wave current injections of 100 pA were elicited serially from −400 to +900, repeated 3 times, with a 3-5-min rest period between each current-injection round, and trials averaged within a single cell. Series resistance was monitored at the beginning and end of each trial. Resting membrane potential was obtained at the initial starting potential in the 100ms prior to the start of current injection. To avoid contamination by hyperpolarization-activated cation currents (I_h_ or “sag”), which were evident in the first 3 hyperpolarizing current steps (−400pA, −300pA, −200pA), input resistance (IR) was measured at the −100pA hyperpolarizing current step. Rheobase was measured as the minimum current required to evoked one or more action potentials during an 800ms depolarizing current step; sag potential was measured as the difference between the maximum negative peak at −400pA and when the voltage reaches steady state. Action potential (AP) number was calculated at each depolarizing current step and AP threshold was measured from the first AP fired. AP accommodation was calculated as the instantaneous frequency at each spike interval. AP waveform properties including AP amplitude, half-width, and after hyperpolarization (AHP) were measured. AP amplitude and AHP were measured relative to threshold and half-width was measured at 50% max amplitude. Instantaneous frequency of firing was measured as the frequency between two consecutive spikes and was plotted against spike interval number. This allows for the measurement of burst like firing as well as accommodation. AP interval number was highly variable among cells making interpretation of accommodation at later spike interval numbers difficult. Therefore, only cells that spiked more than 5 times at +100pA, or more than 17 times at +200pA were used for analysis. Delay to the first AP was measured as the time it took from the start of current injection to the peak amplitude of the first action potential. Signals were collected using an Axopatch 200B amplifier (Molecular Devices, Sunnyvale, CA). Recordings were lowpass filtered at 5kHz, gain 5x, and sampled at 10kHz using a Digidata 1440A (Molecular Devices, LLC, San Jose, CA) and stored on a computer equipped with pClamp10 (Molecular Devices, LLC, Sunnyvale, CA).

### Data Analysis

All data were obtained using the electrophysiology data acquisition software pClamp10 (Molecular Devices, LLC, Sunnyvale, CA.) and analyzed using Origin 2016 (OriginLab), Graphpad Prism 7 (GraphPad Software, Inc.) and SPSS 22 (IBM Corp.). *N* (number) is reported as number of cells with a minimum of 4 animals used per genotype. Experimenter was blind to genotype during experimental procedure, and data collection, with genotype only revealed at final analysis. Results reported at mean ± SEM with significance set at p < 0.05 (*) determined by unpaired Student’s t-test assuming (with Welch’s correction for unequal variance) or Mann-Whitney U-test for nonparametric data. Outliers were determined with a Grubb’s test (GraphPad Software, Inc.) and significant outliers were removed.

## Results

### Excitability is increased in dentate granule cells from TgF344-AD rats

Increased dentate granule cell excitability has been reported in other transgenic rodent models of AD (Hazra et al., 2013; Nenov et al., 2015). We recently reported increased steady-state postsynaptic depolarization of DGCs during high frequency stimulation and heightened LTP magnitude at MPP-DGC synapses in 6 month old TgF344-AD rats (Smith and McMahon, 2018; Goodman et al., 2020). The increased steady-state depolarization is partially driven by heightened activity of β adrenergic receptors (β-ARs) at MPP-DGC synapses, as a β-AR antagonist does not completely eliminate the increased depolarization (Goodman et al., 2020). Therefore, we wondered if heightened intrinsic excitability could also be occurring and tested this using whole-cell current clamp recordings of DGC neurons. Intrinsic excitability was isolated from synaptic transmission using the AMPAR antagonist DNQX (10μM), NMDAR antagonist d,l-APV (100μM) and the GABAergic antagonist picrotoxin (100 μM). Voltage-gated calcium channels Ca_V1.1-4_ were blocked with nifedipine (10μM) to eliminate any contribution from the heightened β-AR function in TgF344-AD rats (Goodman et al., 2020) (**Fig. 1A**). Incremental hyperpolarizing and depolarizing current steps of 100pA, 800ms in duration, were used to assess passive and active membrane properties (**Fig. 1D**). Resting membrane potential (RMP) was measured immediately after obtaining whole-cell configuration in the absence of a current injection (I=0). The average RMP recorded from DGCs in TgF344-AD rats was not different from that measured in Wt littermates (**Fig. 1B**; *t*_(28.73)_ = 1.69, *p* = 0.10) but Wt DGCs were over 4-fold more likely to rest below −80mV than their TgF344-AD counterparts (Wt: 7/22 [31.81%] vs Tg: 1/13 [0.077%]). Input resistance (R_I_) was calculated from the steady-state hyperpolarization generated at −100pA (in which sag current is undetectable). Input resistance in TgF344-AD DGCs was significantly higher than that measured in Wt littermates (**Fig. 1C**; Tg (*Mdn* = 168.1), Wt (*Mdn* = 133.8), *U* = 69.0, *p* = 0.011). Together these data demonstrate modified passive membrane properties in DGCs from TgF344-AD rats.

**Figure 1.**
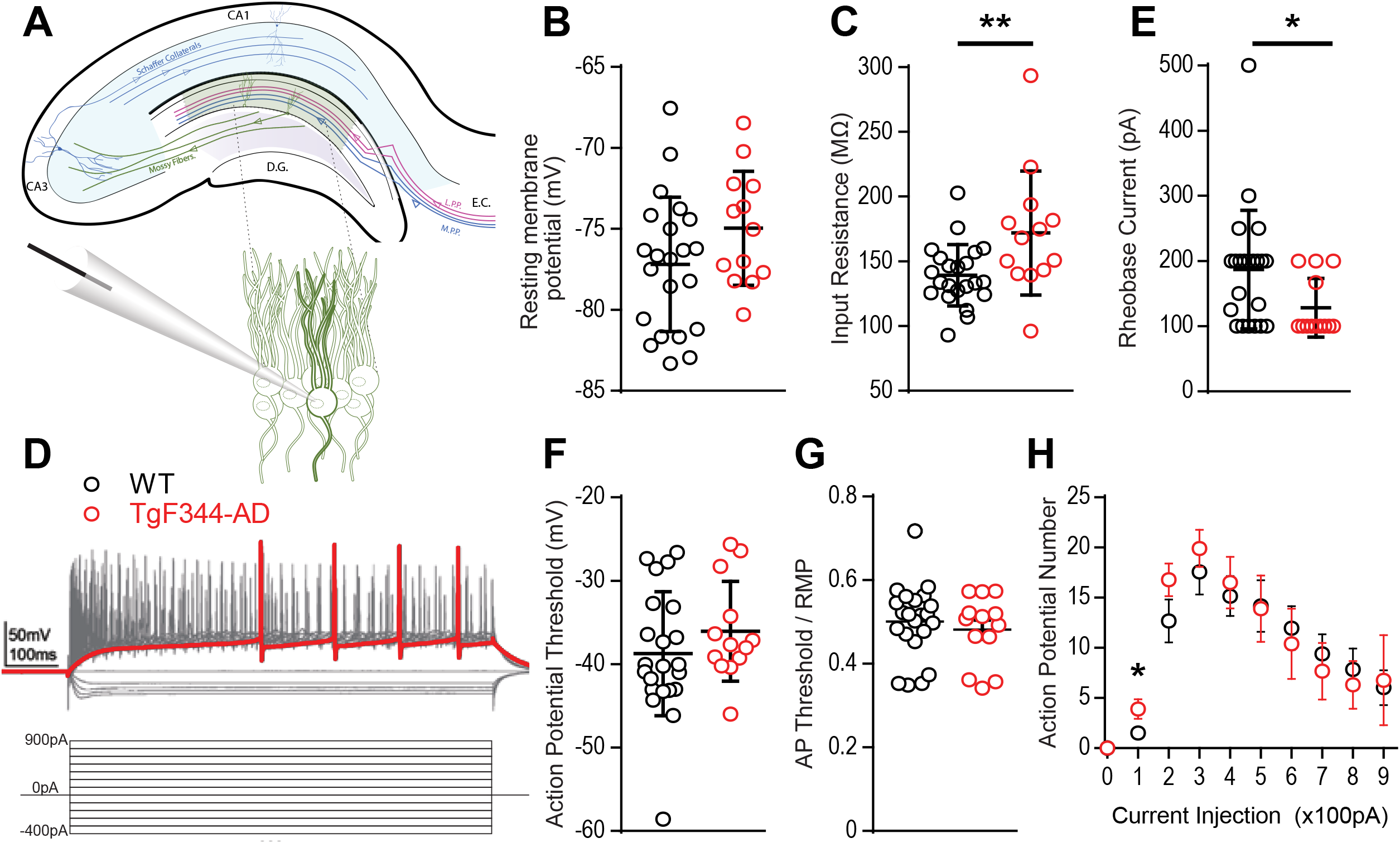
Membrane properties are altered in TgF344-AD DGCs. (A) Schematic of the trisynaptic circuit in a coronal slice from rat hippocampus (top) with expanded recording setup (bottom). (B) Mean resting membrane potential is not different between Wt (black, 22 cells/8 animals) and TgF344-AD females (red, 13 cells/6 animals; p > 0.05, but a higher fraction in Wt have values below −80mV). (C) Input resistance is significantly decreased in TgF344-AD DGCs (Wt n = 22 cells/8animals; Tg n = 13 cells/6 animals; **p ≤ 0.01). Data represent mean ± SEM. Significance determined by unpaired *t*-test. (D) Example traces and current-step protocol (14 serial 100pA hyperpolarizing to depolarizing current steps at 800ms duration from −400pA to +900pA). The red trace represents the first depolarizing current step to fire and action potential. (E) The first depolarizing current step to elicit and action potential, Rheobase (pA), is significantly decreased in TgF344 compared to Wt (*p < 0.05). (F) Action potential threshold is not different between genotypes (p > 0.05). (G) When AP threshold is normalized to RMP there remains no difference between genotype (p > 0.05). (H) Action potential number was enhanced for TgF344-AD rat DGCs at 100pA current injection (*p* < 0.05), but not at higher current injections. For all panels, Wt n = 22 cells/8animals; Tg n = 13 cells/6 animals). Data represent mean ± SEM. Significance determined by unpaired Student’s *t*-test.

### DGCs fire at lower depolarizing membrane potentials in TgF344 rats

Fast voltage gated and slow persistent Na^+^ currents dictate active membrane properties and alterations in both types of Na^+^ currents have been implicated in AD (Verret et al., 2012; Corbett et al., 2013). To determine if active membrane properties are modified in TgF344-AD rats, we measured; action potential threshold, rheobase, and action potential number. Incremental 100pA depolarizing current steps reliably elicited action potentials in DGCs from both TgF344-AD rats and Wt littermates (**Fig. 1D,E**). We did not detect a difference in AP threshold between genotypes when measured from baseline (**Fig. 1F**; *t*_(33)_ = 1.11, *p* = 0.27) nor when AP threshold was normalized to the cell’s RMP (*t*_(33)_ = 0.65, *p* = 0.52) (**Fig. 1G**). Interestingly, the rheobase, or the minimum depolarizing current needed generate an AP, was significantly lower in TgF344-AD rat compared to Wt DGCs (**Fig. 1E**; *t*_(32.27)_ = 2.542, *p* = 0.016). This reduced depolarizing current to generate an AP in the TgF344-AD rat DGC is congruent with the increased input resistance in TgF344-AD DGCs (**Fig. 1E,C**). A decrease in rheobase could lead to an increase in AP number, which was evaluated next at each of the depolarizing current steps. While there was no difference in AP number at current step values higher than 200pA, the TgF344-AD rat DGCs fired more APs at 100pA (**Fig. 1H**; *t*_(25)_ = 2.26, *p* = 0.033). The greater number of APs at this low current step suggest an enhanced sensitivity to fire following a reduced stimulation.

To further investigate initial firing properties, we reasoned that in addition to firing more APs at the minimal depolarizing step as mentioned above, they should also fire more reliably at this minimal depolarizing current injection than Wt DGCs. Indeed, we found that at a depolarizing current injection of +100pA, the probability of successful AP generation is greater in DGCs from TgF344-AD compared to Wt rats, with “0” denoting failure to fire at least one action potential and “1” indicating successful AP generation (**Fig. 2A,B**). In fact, we find that DGCs in TgF344-AD rats were almost two-fold more likely to fire an AP than DGCs from Wt rats at the smallest depolarizing current injection (**Fig. 2C**). Consistent with this observation, the voltage response during the first depolarizing current step (+100pA) in TgF344-AD DGCs is significantly increased (**Fig. 2D**; *t*_(31.27)_ = 3.146, *p* = 0.004), and the delay to fire the first AP was significantly reduced **(Fig. 2E,F**; Tg (*Mdn* = 0.15), Wt (*Mdn* = 0.23), *U* = 9.0, *p* = 0.016). Together these data show that for the same amount of excitable input, TgF344-AD DGCs fire more consistently and experience a greater steady-state membrane depolarization compared to Wt, consistent with the observation that TgF344-AD dentate is more excitable as early as 6 months.

**Figure 2.**
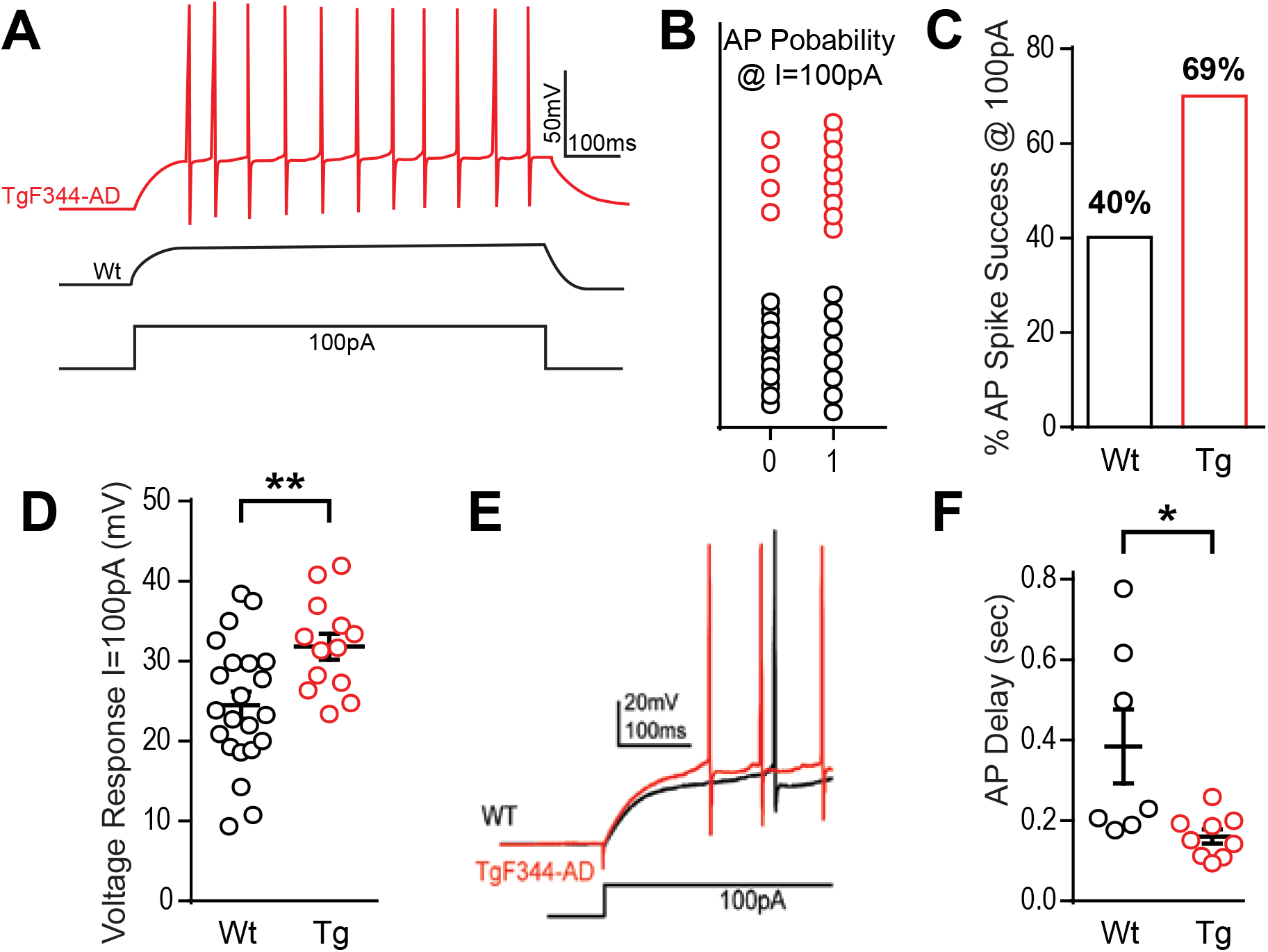
TgF344-AD rats have a lower excitation threshold and fire sooner. (A) Representative traces of the average response of DGCs from TgF344-AD rats vs Wt littermates at the first depolarizing step, +100pA. (B) The probability of firing an action potential at +100pA (0 = did not fire an AP, 1 = fired at least one AP) is greater in TgF344-AD DGCs (Wt 9/22 cells fired, Tg 9/13 cells fired). (C) DGCs from Wt rats discharged at +100pA 40.9% of the time compared to a 69.2% success rate observed in TgF344-AD rats. (D) Regardless of whether the cell discharged or not, TgF344-AD DGC membranes have a greater voltage change (mV) compared to Wt during the first at +100pA (Wt n = 22 cells/8animals; Tg n = 13 cells/6 animals; **p < 0.01). (E) Representative traces of the delay to fire the first action potential (Wt = black, Tg = red). (F) TgF344-AD DGCs had a decreased delay to fire during the first depolarizing current step at +100pA (Wt n = 7 cells/5 animals, Tg n = 9 cells/6 animals; *p < 0.05). Data represent mean ± SEM. Significance determined by unpaired Student’s *t*-test.

### Action potential kinetics do not account for increased excitability in TgF344-AD DGCs

To assess whether changes in rheobase and the increased depolarizing voltage response (**Figs. 1E, 2D)** impact the AP kinetics, we quantified several properties of the AP waveform to include amplitude, half-width, and the amplitude of afterhyperpolarization (AHP) (**Fig. 3A**). We chose to quantify kinetics of the first AP fired at rheobase, to prevent modification of AP waveform due to inactivation of Na^+^ and K^+^ channels, as well as activation of currents at hyperpolarized potentials. The first AP waveform was averaged from all cells recorded in each group (Wt = black, Tg = red) (**Fig. 3B**). AP amplitude (**Fig. 3C**), half-width (**Fig. 3D**), and AHP amplitude (**Fig. 3E**) were not significantly different between genotypes (p > 0.05). Together these data suggest voltage gated Na^+^ and K^+^ channels that mediate membrane depolarization and repolarization are not functionally altered at this early pathological stage in TgF344-AD rats.

### Sag-mediated current is decreased in TgF344-AD DGCs

**Figure 3.**
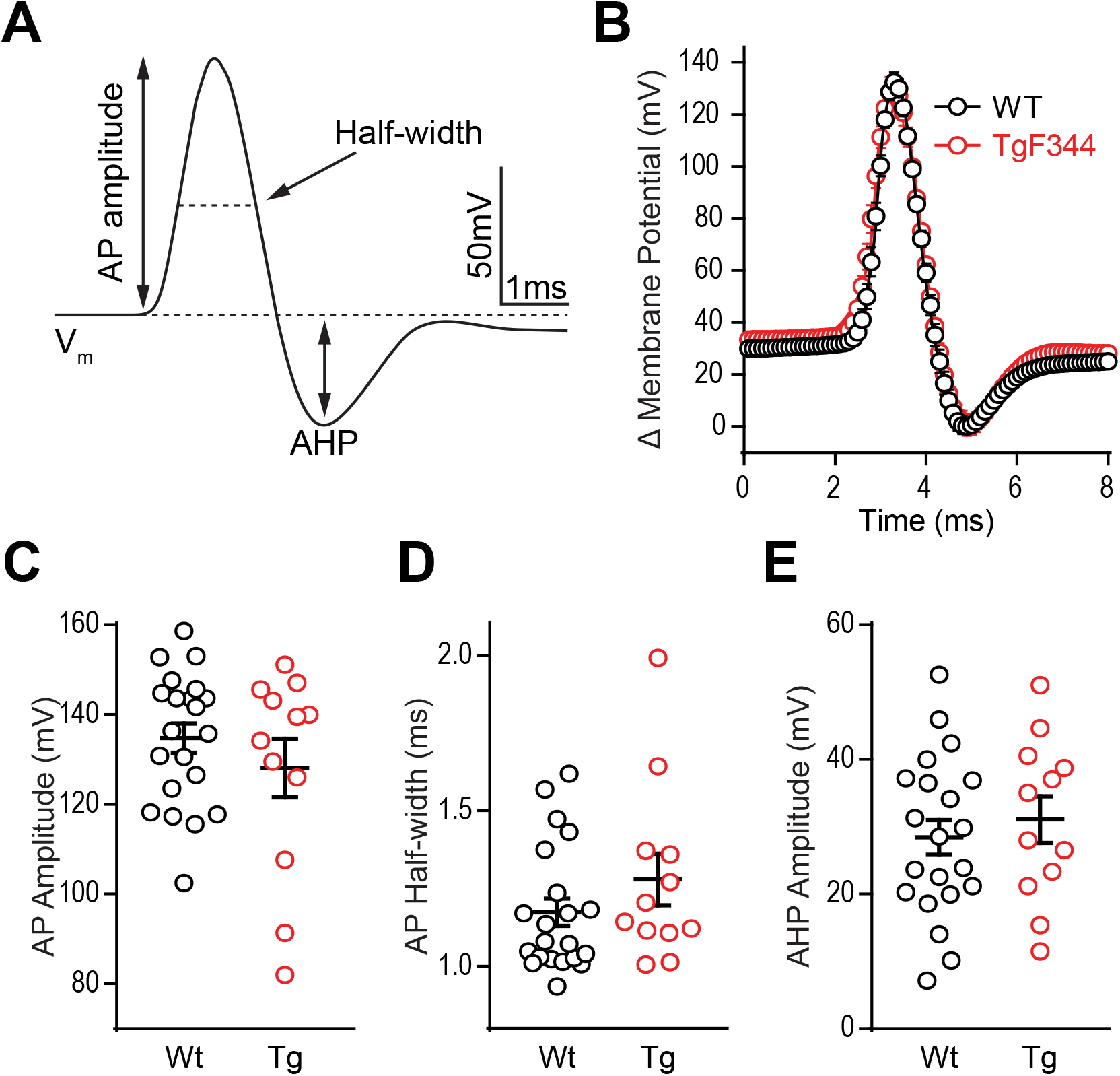
Action potential waveform properties are not different between TgF344-AD and Wt DGCs. (A) Schematic of measurements derived from a typical action potential waveform including, AP amplitude, half-width, and afterhyperpolarization (AHP). (B) Averages of the first action potential fired at rheobase in each genotype (error bars represent SEM). (C) Action potential height (mV), (D) action potential half-width (ms), and (E) AHP amplitude (mV) in DGCs from 6 month TgF344-AD and Wt female rats are not different. For all panels Wt n = 22 cells/8animals, Tg n = 13 cells/6 animals, p > 0.05. Data represent mean ± SEM. Significance determined by unpaired Student’s *t*-test.

Hyperpolarization activates hyperpolarization-activated-cyclic-nucleotide gated currents, I_h_ (HCN channels), which mediate excitability and rhythmic firing in neurons. DGCs typically contain little I_h_ current, yet in patients with severe epilepsy, increased hippocampal HCN channel expression is confined to dentate (Bender et al., 2005; Poolos et al., 2002; Poolos and Johnston, 2012), and may represent a compensatory mechanism in response to over-excitation. To determine if decreased HCN channel function could explain the increased excitability observed at this early stage, we additionally measured the “sag”, a voltage signature of HCN channels (**Fig. 4A**). At −400pA, DGCs in TgF344-AD rats did not show a significant decrease in sag amplitude (**Fig. 4B**; *t*_(21.43)_ = 1.181, *p* = 0.25), but we noted an enhanced voltage response in DGCs from TgF344-AD rats at the −400pA current injection used to measure sag (**Fig. 4C**; *t*_(18.44)_ = 2.27, *p* = 0.04), which is consistent with the observation that R_I_ is increased in TgF344-AD rats. To compensate for the enhanced R_I_, sag amplitude was normalized to the hyperpolarized voltage response which resulted in a significant decrease in sag amplitude in DGCs fromTgF344-AD rats DGC sag amplitude (**Fig. 4D**; *t*_(29.94)_ = 2.09, *p* = 0.045). When taken together, the reduced sag/voltage response may indicate that I_h_ current mediated by HCN channels is impaired in DGCs of TgF344-AD rats.

**Figure 4.**
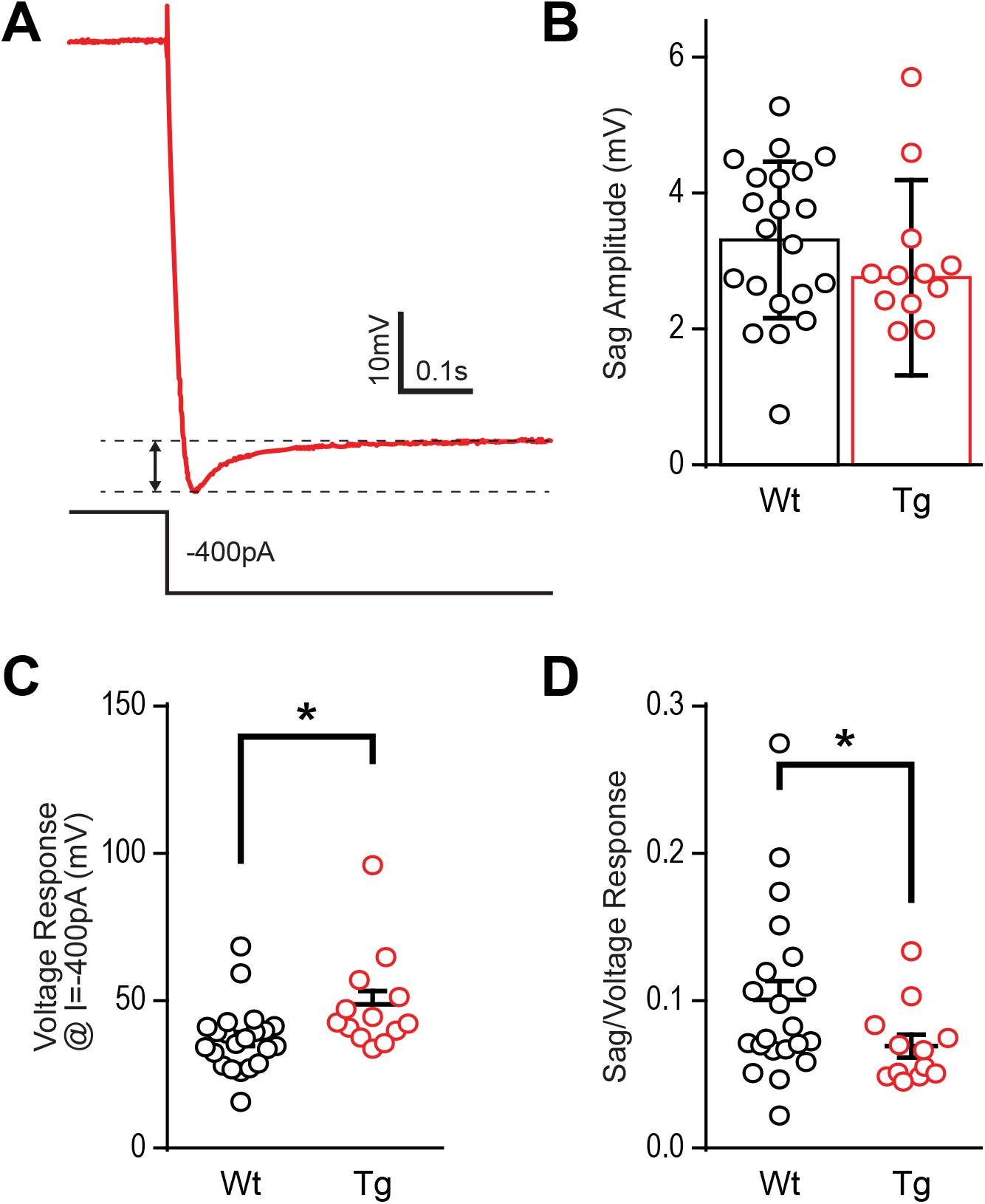
Sag is decreased in TgF344-AD DGCs. (A) Example trace of ‘sag’ current measurements taken at −400pA current injection. (B) The raw amplitude of the sag current is not different between TgF344-AD and Wt littermate DGCs (Wt n = 21 cells/8animals; Tg n = 12 cells/6 animals; *p* > 0.05). (C) The voltage response at −400pA is greater in TgF344-AD rats (Wt n = 22 cells/8animals; Tg n = 13 cells/6 animals; \**p* < 0.05). (D) When Sag amplitude is normalized to voltage response a significant deficit in TgF344-AD DGC sag is unmasked (Wt n = 21 cells/8animals; Tg n = 12 cells/6 animals; \**p* < 0.05). Data represent mean ± SEM. Significance determined by unpaired Student’s *t*-test.

### TgF344-AD DGCs have increased initial spike frequency and greater accommodation

During prolonged depolarization, neurons undergo spike frequency accommodation, where a reduction in AP frequency occurs over time (Madison and Nicoll, 1984) and is mediated by gradual activation and recruitment of K^+^ channel currents. Enhanced excitability of DGCs in TgF344-AD rats may be a result of increased input resistance and reduced rheobase, yet there is no increase in AP number above 100pA current injection suggesting that accommodation may clamp AP number at higher currents. At the first depolarizing step, +100pA (**Fig. 5A**), differences in instantaneous firing frequency are not interpretable since overall spike number is low and several cells fail to spike. To address this, we assessed accommodation at +200pA (**Fig. 5C, D**) when 92% of cells fire an action potential as in (Brown et al., 2011). We found that DGCs in TgF344-AD rats begin with a mean instantaneous firing frequency of 76.15 ± 6.33Hz and accommodate to 20.52 ± 2.07Hz by 17 spike intervals (**Fig. 5B**, red). In contrast, Wt DGCs have a lower initial spike frequency (59.05 ± 9.85Hz) and accommodate to 31.42 ± 3.71Hz by steady state (**Fig. 5B**, black). The initial instantaneous firing frequency is not different between TgF344-AD DGCs compared to Wt (**Fig. 5B**; p > 0.05). The second and third intervals, however, suggest a greater instantaneous firing frequency for Tg DGCs compared to their Wt littermates (*t*_(22.70)_ = 2.032, *p* = 0.054; *t*_(22.68)_ = 2.275, *p* = 0.032, respectively). When the final 5 spike intervals (15-20) are collapsed (**Fig. 5B**, dashed box), the TgF344-AD rat DGCs have a significantly reduced instantaneous frequency (*t*_(9.66)_ = 11.08, \*\*\*\**p* < 0.0001), indicating a greater accommodation. Together these data suggest TgF344-AD DGCs accommodate to a greater degree than Wt, but that the accommodation mechanism is slower to engage. To further validate this finding a larger depolarizing step (+300pA) was also used to measure accommodation with similar results as in (Brown et al., 2011) (**Fig. 5C**). The first spike interval is significantly shorter for TgF344-AD rat DGCs (*t*_(23.54)_ = 2.906, *p* = 0.0078) but quickly becomes comparable to Wt at the second and third interval (*t*_(29.93)_ = 1.758, *p* = 0.089; *t*_(26.31)_ = 0.677, *p* = 0.504, respectively) (**Fig. 5C**). When the final 7 spike intervals (19-25) are collapsed (Fig. 5C inset), the TgF344-AD rat DGCs have a significantly reduced instantaneous frequency (*t*_(11.98)_ = 20.13, \*\*\*\**p* < 0.0001), again indicating a heightened accommodation. These findings are consistent with the interpretation that the lack of change in total AP number at a given depolarizing current step is a consequence of a simultaneous increase in instantaneous AP frequency and a greater frequency accommodation in DGCs of TgF344-AD rats.

**Figure 5.**
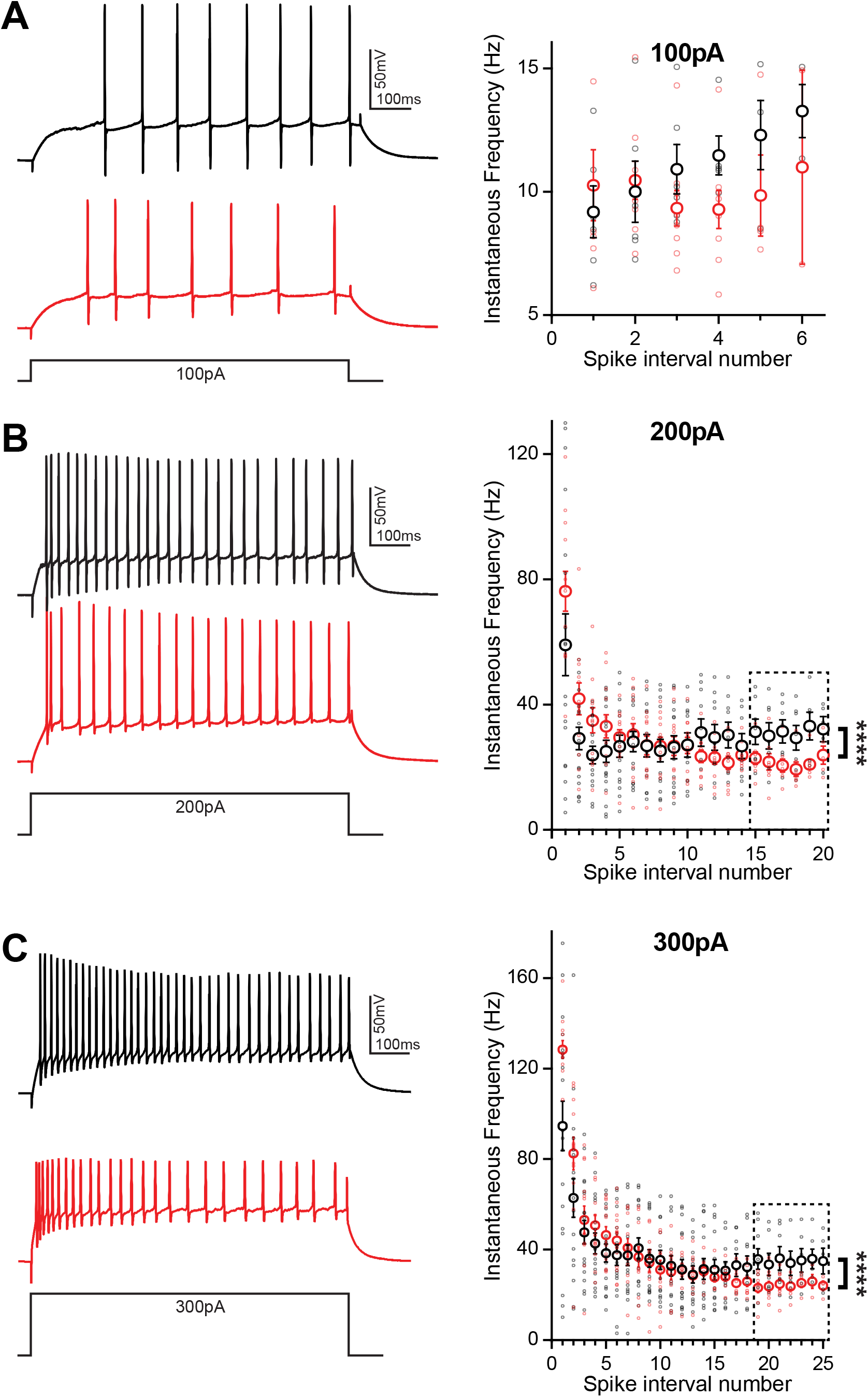
Initial spike frequency and spike accommodation rate are greater in TgF344-AD DGCs. (A) (left) Graph of instantaneous firing frequency (Hz) between action potentials (spike intervals) at 100pA to include TgF344-AD (red) and Wt (black) DGCs that successfully fired an action potential at +100pA current injection. (right) Example traces. (B) (left) Graph of instantaneous firing frequency (Hz) between action potentials (spike intervals) at 200pA showing both increased initial spike frequency and greater accommodation (inset) (*p*<0.0001). (right) Example traces. (Wt n = 7 cells/5 animals; Tg n = 5 cells/5 animals)(C) (left) Graph of instantaneous firing frequency (Hz) between action potentials (spike intervals) at 300pA showing both increased initial spike frequency and greater accommodation (inset) (*p*<0.0001). (right) Example traces. (****p < 0.0001 Wt n = 7 cells/5 animals; Tg n = 7 cells/5 animals). Data represent mean ± SEM. Significance determined by unpaired *t*-test with Welch’s correction.

## Discussion

Hyperexcitability in hippocampal circuits is reported in many mouse models of AD (Palop et al., 2007; Roberson et al., 2007; Rocher et al., 2008; Verret et al., 2012; Hazra et al., 2013; Šišková et al., 2014; Nenov et al., 2015). However, in transgenic rat models of AD there is a lack of information regarding intrinsic excitability, especially in the dentate gyrus hippocampal subfield. Here we explored whether intrinsic excitability is altered in the novel TgF344-AD rat model, as it might contribute to synaptic changes such as the increased steady-state depolarization during high-frequency tetanus and the enhanced LTP magnitude we previously reported in TgF344-AD rats, (Smith and McMahon, 2018; Goodman et al., 2020), which more faithfully recapitulates human AD pathology than other transgenic rodent models (Do Carmo and Cuello, 2013; Cohen et al., 2013; Rorabaugh et al., 2017).

Passive membrane properties of neurons determine resting membrane potential which can be altered by the presence of soluble Aβ(Fernandez-Perez et al., 2016). We did not detect a genotype difference in resting membrane potential of DGCs, but did find that DGCs in TgF344-AD rats had both a greater likelihood of resting above −80mV and an elevated R_I_ (**Fig. 1C,D**). A depolarizing shift in RMP has been reported in cortical and CA1 pyramidal cells in various transgenic tau-expressing or AD mouse lines (Minkeviciene et al., 2009; Verret et al., 2012). K^+^ leak and G-protein coupled inwardly rectifying K^+^ (GIRK) channels play a major role in determining RMP (Lüscher and Slesinger, 2010) and their membrane expression could be directly linked to measures of IR. Future studies should explore whether changes in these channels underlie the increased R_I_, or even the increased likelihood of a depolarized RMP in the DGCs of TgF344-AD rats.

The enhanced input resistance is further evidenced by the heightened voltage response to both depolarizing and hyperpolarizing currents (**Figs. 1C, 2D**, respectively). Furthermore, the current required to produce APs in TgF344-AD rats is reduced (**Fig. 1E**). Upon closer inspection, the minimal depolarizing current injection used to elicit APs (+100pA) showed both a significant enhancement in the probability of AP firing, and a decreased delay to fire in DGCs from TgF344-AD rats DGCs (**Fig. 3**). It remains unclear whether these changes are due to altered ion channel expression or function.

K^+^ channels function to dampen membrane excitability and impaired K^+^ channel expression and function is implicated as a mechanism for hyperexcitability in AD models (Scala et al., 2015). Specifically, K^+^ channel dysfunction should reduce the AP width and AHP magnitude (Tamagnini et al., 2015). However, we did not detect a decrease in AP width during the falling phase (Fig. 4D), suggesting delayed rectifying K^+^ channels, are intact at this age. The AHP shape is mediated by Ca^2+^-activated K^+^ channels such as the large conductance BK-type channels and the small conductance SK-type channels (Andrade et al., 2012). Unsurprisingly, we did not observe a difference in AHP amplitude in TgF344-AD rats (**Fig. 3E**), suggesting decreased BK and/or SK-type mediated currents likely do not underlie the changes we see. Alternatively, increased intrinsic excitability can be linked to enhanced LTP by an A-type K^+^ channel mediated increase in back propagating APs resulting from enhanced dendritic excitability, and Ca^2+^ influx (Frick et al., 2004).

Membrane repolarization is mediated in large part by I_A_, which dampens excitatory post synaptic potentials (EPSPs), raises the threshold for AP initiation, and suppresses back propagating action potentials (Hoffman et al., 1997). Activation of I_A_ clamps the membrane below threshold thereby determining the delay to the first action potential spike (Storm, 1990). We calculated the time to first spike in DGCs from TgF344-AD rats and Wt littermates a revealed a decreased delay to fire an AP in TgF344-AD rats (**Fig. 2E**), suggesting the effect of I_A_ on dampening excitability may be compromised. While our somatic recordings did not reveal a difference in AHP, whether the threshold for local dendritic I_A_ current is altered, or if a down-regulation of K^+^ current is responsible for the increased excitability observed here awaits additional investigation.

Voltage gated Na^+^ channels (VGNaC) are responsible for the initiation of APs and tightly control AP threshold. Decreased VGNaCs have been previously observed in AD-mouse models (Verret et al., 2012; Corbett et al., 2013). Specifically, the voltage gated sodium channel, Na_V1.1_, is decreased in glutamatergic and GABAergic neurons in Tg-AD mouse models leading to increased hyperexcitability, likely through enhanced E/I ratio (Verret et al., 2012; Corbett et al., 2013). We find AP threshold is not different between TgF344-AD and Wt (**Fig. 1F,G**). Furthermore, changes to VGNaC should produce changes to AP amplitude or half-width, yet we did not detect a difference in either measure (**Fig. 3C,D**). Together, these data suggest VGNaC function on DGCs is intact in TgF344-AD rats at 6-8 months, yet whether these channels are decreased on interneurons that innervate DGCs remains to be investigated (Palop et al., 2007; Verret et al., 2012). Unlike the rapidly activating/inactivating VGNaC, persistent Na (I_NaP_) currents do not inactivate, lasting for hundreds of milliseconds with the ability to influence rheobase. These I_NaP_ can augment cell excitability with an additive effect to other depolarizing currents experienced by the cell, can reduce rheobase, and have been implicated in epileptic firing. Our data show a significant decrease in rheobase (**Fig. 1E**) which may indicate a change in I_NaP_.

Hyperpolarization activated currents can mediate DGC excitability and HCN channel function is impaired in Tg-mouse models of AD (Kaczorowski et al., 2011). While the raw amplitude of the sag current was not different, we found enhanced hyperpolarized shift in the voltage response in TgF344-AD rat DGCs (**Fig. 4C**). Importantly, when the sag amplitude is normalized to voltage response, a deficit in sag response is unmasked, suggesting that for the same amount of membrane hyperpolarization, fewer HCN channels are activated (**Fig. 4D**). These data support that decreased HCN channel function could mediate the DGC hyperexcitability we report in TgF344-AD rats at 6 months.

Interestingly, while the overall number of APs was not different between genotypes, both the initial AP frequency and spike accommodation were elevated in TgF344-AD DGCs, suggesting the dynamics of AP firing frequency are also altered in TgF344-AD rats. AP number was highly variable among cells in each group, and therefore cells that did not meet a minimum spike number (5 spikes at 100pA, or 17 spikes at 200pA and above) were excluded for accommodation analysis. Interestingly, when these low-fidelity cells were entered into the analysis, the difference in initial firing frequency was abolished, regardless of spike number, and therefore we cannot rule out the possibility that we were recording from two populations of mature dentate granule cells (Nenov et al., 2015). In fact, variability in spike number among cells even within the same group indicate there may be multiple populations of mature DGCs from which we are recording (Nenov et al., 2015). While we did not directly measure DGC bursting activity, enhanced instantaneous frequency within the bursting range of 3-8 APs in the TgF344-AD rat may enhance the propagation of signals through the dentate or functionally rearrange feedforward inhibition (Neubrandt et al., 2018). This may be especially true of signals which otherwise would not evoke a DGC AP due to their increased R_I_. Changes in initial firing frequency and spike rate accommodation support a role for altered Na^+^ and K^+^ channel function in DGCs during short or extended depolarization and therefore future studies should aim to determine their functional role in this AD model.

## Conclusions

Excitation inhibition imbalance is an early feature of pathology in the hippocampus of preclinical AD patients, and this imbalance has been recapitulated in several models of AD-like pathology. We previously reported increased steady state depolarization and LTP magnitude in TgF344-AD DG at 6 months. While the enhanced LTP was dependent on heightened β-AR function, this enhanced function was not sufficient to account for the heightened SSD (Smith and McMahon, 2018; Goodman et al., 2020). Here we used whole-cell current clamp to show that intrinsic excitability is increased in DGCs from TgF344-AD rats and provides one mechanistic explanation for synaptic alterations and may contribute to early increase in LTP magnitude reported in this model.

